# The potent anti-influenza virus activity of FluPep is mediated by interactions with cellular anionic polysaccharides

**DOI:** 10.64898/2026.09.15.751766

**Authors:** Zaid K. Alghrair, Bahram Ebrahimi, David G. Fernig

## Abstract

“FluPep” is a 16 amino-acid peptide with potent anti-viral activity against influenza A virus, including the H1N1 subtype. To understand how FluPep exerts its effects, gold nanoparticles were functionalised with FluPep, since gold nanoparticles provide a very sensitive probe that can be detected optically. Our results demonstrate that nanoparticle-FluPep conjugates did not bind to influenza virus particles *in vitro*. However, the nanoparticle-FluPep conjugates bound to polyanions on the cell surface, and after washing cells they remained cell-associated and were able to inhibit ‘flu virus infectivity. Nanoparticle-FluPep conjugates bound to heparin affinity columns and their binding to cells could be reduced by pre-treatment of the cells with heparinase and neuraminidase, but not with chondroitinase ABC. These data indicated that FluPep bound to heparan sulfate and sialic acid-containing glycans in the pericellular matrix and in doing so prevented viral infectivity. The importance of the interaction of FluPep with heparan sulfate was examined by extending FluPep at its C-terminus with amino acid sequences responsible for the interactions of growth factors with heparin. The resultant two new engineered peptides, termed ‘superFluPep1’ and ‘superFluPep2’, were more potent than parental FluPep. The results demonstrate that FluPep likely exerts its antiviral activity by interfering with virus-cell interactions and that enhancing the binding of such peptides to heparan sulfate is an effective strategy to increase their anti viral potency.

## Introduction

Influenza virus or the ‘flu’ virus is a highly infectious disease for both human and farm animals caused by RNA viruses belonging to the *Orthomyxoviridae* family (Chavas et al., 2010). The flu virus is mainly transmitted by aerosols and infects the upper respiratory tract (Cowling et al., 2013). The infection cycle is considered to be initiated by the interaction of the viral heamagglutinin (HA) protein with sialic acids on cell surface glycoproteins (Das et al., 2010). Following endocytosis of the viral particle, the reduced pH within the endosomes results in the fusion of the viral envelope and endosomal membranes and the eventual transfer of the viral RNA genome segments into the cell nucleus (Brunotte et al., 2014). These steps launch viral genome replication, synthesis of viral proteins and so the generation of new viral particles, which then bud from the infected cell (Stubbs and te Velthuis, 2014) (Leser and Lamb, 2005). The virus binding and membrane fusion, replication and budding of virions have been targets for the development of anti-influenza drugs.

There are now a number of peptides with potent anti-flu activities, such as Cyanovirin-N (CV-N) (O’Keefe et al., 2003), a 20-amino-acid peptide (EB, for entry blocker) derived from the signal sequence of fibroblast growth factor 4 (Jones et al., 2006), and Flufirvitide (Skalickova et al., 2015). Other classes of molecules have also been shown to inhibit influenza virus infectivity, such as sialic acid derivatives (Matsubara et al., 2010) and sulfated polysaccharides, including the glycosaminoglycan heparin (Skidmore et al., 2015), (Baba et al., 1988), likely by preventing virus binding, though the mechanism has yet to be defined. The peptide FluPep inhibits infectivity of influenza A virus, including the H1N1 subtype, in cultured cells and in mouse models with very high potency (Nicol et al., 2012). FluPep was considered to exert its antiviral activity from the outside of the cell, by binding to HA on virus particles (Nicol et al., 2012). However, the latter measurements were indirect, due to the difficulty in measuring the binding of a peptide, so the target of the FluPep peptide has remained inconclusive. We demonstrated that FluPep conjugated to gold and silver nanoparticles (AuNPs and AgNPs, respectively) retains its anti-flu activity and indeed this is enhanced at low stoichiometries of conjugation (Alghrair et al., 2019). Noble metal nanoparticles provide a very sensitive probe that can be detected optically. This is due to their surface plasmons, which make these materials the strongest absorbers and scatters of light (Haiss et al., 2007, Paramelle et al., 2014). We have, therefore, taken advantage of the optical properties of AuNP and used AuNP-FluPep conjugates to determine whether FluPep binds to virus or to the cell, as a means to start to decipher the anti-flu mechanism of the FluPep. The results show that FluPep does not interact with the virus particle but instead interacts with two anionic components of the cell surface/pericellular matrix. FluPep functionalised AuNP bound to heparin affinity columns, whereas digestion of cells with chondroitinase ABC, heparinase and sialidase demonstrated that FluPep-AuNP conjugates interacted with heparan sulfate (HS) and sialic acid containing glycans. By adding additional amino acid sequences derived from heparin-binding sites of growth factors to target FluPep to HS of the pericellular matrix we demonstrated that this strategy enhanced further the anti-viral activity of FluPep.

## Materials & methods

### Materials

#### Peptides and AuNP

FluPep (Nicol et al., 2012),derived peptides and the peptidol ligand (Table 1) were purchased from Peptide Protein Research (PPR Ltd, Hampshire, UK). The alkanethiol ethylene glycol ligand, HS-EC_11_-EG_4_, was purchased from Prochimia (ProChimia Surface Sp. z o.o., Sopot, Poland). AuNP of 8.8 nm diameter and stabilized in citrate buffer were purchased from British Biocell (BBInternational Ltd, UK). Nanosep filters, 10 kDa and 100 kDa cut off, were from PALL (PALL Corp., Portsmouth, Hants, UK). UV-vis spectra (2 nm incremental steps) were measured using a SpectraMax Plus spectrophotometer (Molecular Devices, Wokingham, UK), 384-well plates were from Corning (Lowell, US) and the concentration of AuNP was determined at 450 nm (Haiss et al., 2007).

**Table 1:**
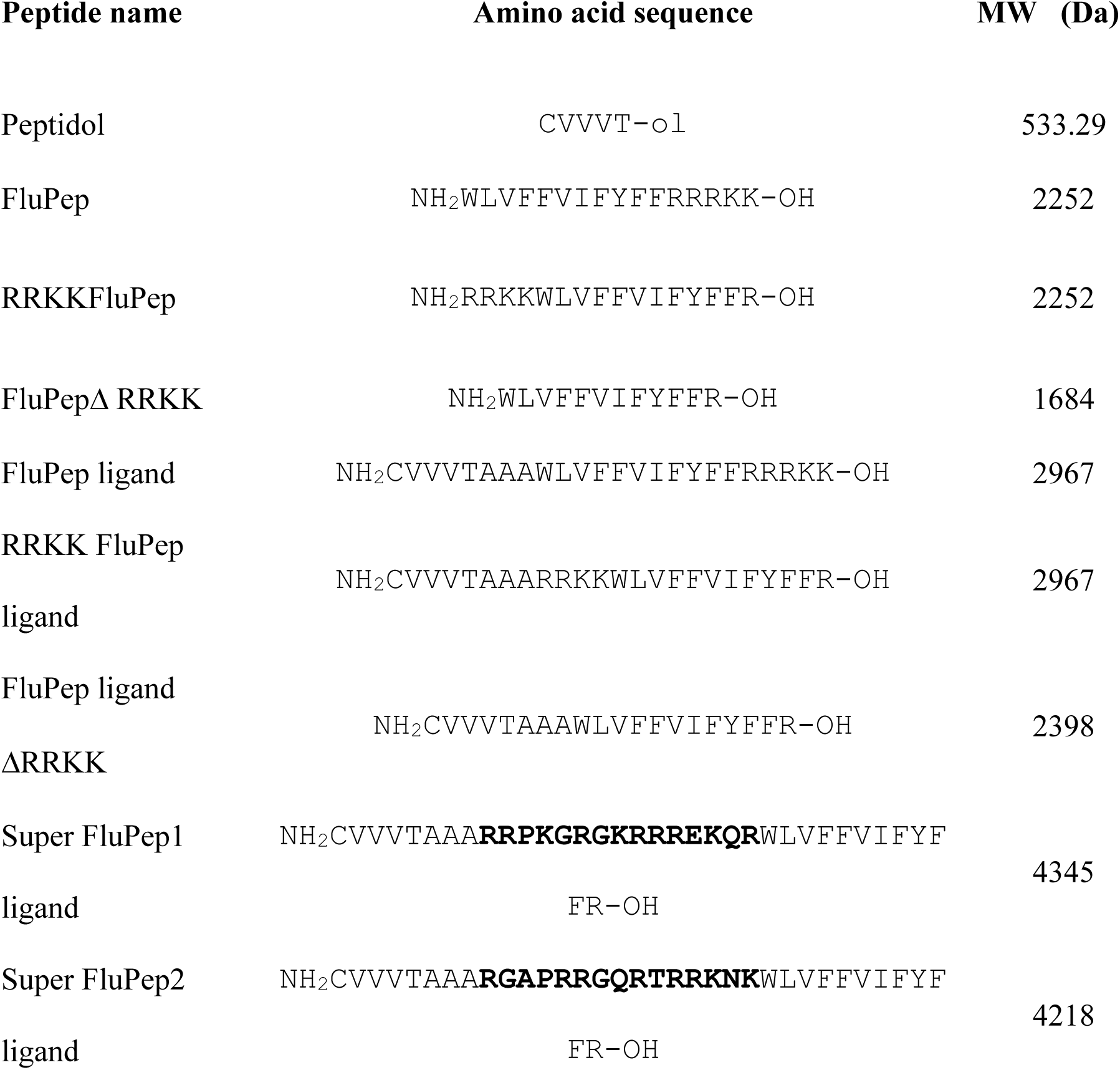
Peptide names, amino acid sequences and their molecular weight. “Ligand” is appended to the peptide name when it contains the N-terminal sequence CVVVT necessary for conjugation to AuNP. RRKKFlu Pep has its C-terminal basic sequence placed at the N-terminus. A spacer of two alanine residues was incorporated between the CVVVT ligand sequence and the FluPep sequences. Super FluPep1 has a sequence (in **bold**) replacing the RRKK sequence of RRKKFluPep. This sequence was derived from placental growth factor2 (PLGF2) and corresponded to a sequence used previously to enhance growth factor HS binding (Martino et al., 2014). Super FluPep2 has a sequence (in **bold**) replacing the RRKK sequence of RRKKFluPep derived from to the canonical heparan binding site of fibroblast growth factor10, (FGF10) (Xu et al., 2012). To enhance its HS binding further, the N-terminal K was replaced with the R from the corresponding sequence of FGF22 and the C-terminal T with the corresponding K from FGF7 (Xu et al., 2012) (Li et al., 2016)

#### Synthesis of FluPep functionalised AuNP

Mix matrix ligands 70:30 (mole:mole) CVVVT-ol: HS-(CH2)_11_-EG_4_-OH were prepared as described (Duchesne et al., 2008a) by first diluting 35 µL CVVVT-ol (4 mM DMSO:H_2_O) with 35 µL ddH_2_O and 6 µL HS-C_11_-EG_4_-OH (2 mM) with 6 µL EtOH and 18 µL ddH_2_O. Adding the two solutions together yielded a 2 mM ligand solution of 70% (mole/mole) CVVVT-ol and 30% (mole/mole) HS-C_11_-EG_4_-OH. The ligand mixture was added to 900 µL AuNP and mixed by vortex. Then 100 µL of 10x phosphate-buffered saline (PBS: 137 mM NaCl, 3 mM KCl l, 8 mM Na_2_HPO_4_, 15 mM KH_2_PO_4_) with Tween-20 (0.1 % v/v) pH 7.4 was added to AuNPs (Duchesne et al., 2008a) vortexed and the AuNPs placed on a rotating wheel for 24 h. AuNPs were concentrated 10 x by centrifugation on 10 kDa Nanosep centrifugal filters (PALL Corp., Portsmouth, Hants, UK) for 7 min at 10000 rpm (12,000 x g) and the AuNPs diluted with 1x PBST (PBS with 0.05% (v/v) Tween-20). The AuNPs were then further separated from excess ligands by applying (100 µL) to a 5 mL Sephadex G25 gel filtration column with PBS as a mobile phase.

To functionalise the AuNPs with functional peptide ligands, e.g., FluPep ligand (Table 1), these were incorporated into the initial ligand mix at the mole % indicated in the figure legends. However, the separation of these ligands (Mw > 1000 Da) required in addition to gel-filtration on Sephadex G25 six washes, each involving a 10-fold dilution of the AuNPs on a 10 kDa cut off Nanosep filter.

#### Purification of functionalised gold AuNPs

Ion-exchange and heparin affinity chromatography were performed on homemade micro columns of Diethyl-Amino-Ethyl (DEAE) Sepharose, Carboxy-Methyl (CM) Sepharose (both from GE Healthcare) and heparin agarose (BioRad, Hemel Hempstead, UK). The chromatography gel was packed into a white pipette tip (200 µL) using half the filter as a frit at the bottom and equilibrated in PBS. Capped AuNPs were concentrated and exchanged into PBS on a 10 kDa Nanosep centrifugal filter. The AuNPs were then applied to the column and the unbound fraction was recovered. Columns were washed with PBS and eluted stepwise with 1 M NaCl and 2 M NaCl in PBS pH 7.2. The relation of bound and unbound AuNPs to the mole % of functional ligand in the original ligand mixture was analysed based on quantification of the AuNPs. The data indicate that at 0.03% (mole/mole) functional ligand, 10 % of the AuNPs bound the column and thus most (∼95 %) of these AuNPs will possess a single functional ligand, as determined previously (Levy et al., 2006). At higher mole % the number of functional ligands per AuNP will increase. Thus, the purification of the functionalised AuNPs allowed the effects on influenza virus infectivity of largely mono-*versus* pluri-functionalisation to be determined.

#### Ligand exchange assay and Calculation of the aggregation parameter (AP)

Purified ligand capped AuNPs (57 µl), 33 µl 10X PBS and 10 µl DTT solutions at different concentrations (or milliQ water when the required concentration of DTT was 0 mM) were added to a 384-well plate. A blank well, which only contained 100 µl milliQ water, was used as reference. Spectra were acquired on duplicate wells at indicated times.

The surface plasmon absorption peak of 8.8 nm diameter AuNPs is at 520 nm. When AuNPs are aggregated, their surface plasmons couple, causing a red shift in their plasmon absorbance to nearly 650 nm. The aggregation parameter (AP) was defined as (A _650 nm_-A_ref650nm_)/ (A_520nm_-A_ref520_), where A_650nm_ and A_520nm_ are the absorbance of AuNPs at 650 nm and 520 nm, respectively, and A_ref650nm_ and A_ref520_ are the absorbance of water at 650 nm and 520 nm, respectively (Chen et al., 2012). For comparison of results, this primary stability parameter was normalised by dividing the AP value of control ligand capped AuNPs measured in milli Q water where [DTT] = 0.

#### Cell culture

Madin-Darby canine kidney epithelial cells (MDCK) were grown as described (Alghrair et al., 2019) (Alghrair et al., 2019) in Dulbecco’s modified Eagle’s medium (DMEM) supplemented with 5% (v/v) heat inactivated foetal calf serum (FCS) (Labtech International Ltd, East Sussex, UK), 1% (v/v) 200 mM L-glutamine, 1% (v/v) 100 U/mL penicillin and 1% (v/v) 100 µg/mL streptomycin (Gibco, Life Technologies, UK) and incubated in a humidified environment at 37°C under 5% (v/v) CO_2_ atmosphere. Cells were passaged when cell density was approximately 70% by washing with 3 ml of Versene and then trypsinisation with 0.05% (w/v) trypsin-Versene (all Gibco, Life Technologies, UK) and plated at a dilution of 1:4.

#### Virus plaque assay

Plaque assays were performed as described (Alghrair et al., 2019)(Alghair et al., 2019). MDCK in 6-well plates (STARLAB international, Hamburg, Germany, 10^6^ cells/well) were grown until a uniform monolayer was formed. The cells were then infected with a serial dilution of influenza virus inoculum, sufficient to obtain approximately 100 plaques per well, for 1 h at 37 °C on a rocking platform for 1 h. The medium was removed and 2 -3 mL of a 1 % (w/v) agarose overlay (equal volumes of 2 % (w/v) of pre-warmed (55°C) low melting agarose solution (Melford Laboratories Ltd, Blideston Road, Ipswich, UK) and the overlay solution (14 mL 10x MEM, 3.7 mL 7.5% (w/v) bovine serum albumin (fraction V, Sigma-Aldrich), 1.4 mL L-glutamine, 2.6 mL 7.5 % (w/v) NaHCO_2,_ 1.4 mL 1 M HEPES, 1.4 mL (1% (v/v) 100 U/mL penicillin and 1% (v/v) 100 µg/mL streptomycin), 44.8 mL H_2_O and 5µL *N*-acetyl trypsin (Sigma-Aldrich)) was quickly added to the wells. This was left to set for 15 min at room temperature, the plates inverted and incubated for 3 days (37°C, 5% (v/v) CO2). The cells were then fixed with 4 mL 10 % (v/v) neutral buffered formalin (Leica Biosystems Peterborough Ltd, Bretton Peterborough, Cambridgeshire) for 1 h, then the formalin and overlay were removed and cells were stained with 0.1% (w/v in water) toluidine blue, rinsed in water, and left to dry and plaques counted.

#### Preparation of influenza virus stock

Virus stocks were prepared as before(Alghrair et al., 2019) (Alghrair et al., 2019). MDCK cells (90% confluent in T25 tissue culture flasks, VWR, Lutterworth, Leicestershire, UK, 7 x 10^6^ cells/flask) were incubated with 2 mL virus (A/WSN/33 H1N1 subtype), multiplicity of infection (MOI) of 0.001 for 1 h at 37 ^0^C on a rocking platform after washing with 2x5mL PBS. Unbound virus was removed and the cells washed with 2 x 5 mL DMEM. After addition of 5 mL N-acetyl trypsin (Sigma-Aldrich, Merck, UK), 2.5 µg/mL in DMEM, cells were incubated 24-48 hours at 37^0^C until a significant cytopathic effect had developed such that the cells were detaching from the culture substrate. Medium was collected and centrifuged for 5 min at 2500 rpm to remove cell debris and the supernatant containing virus stock, was stored at -80^0^C.

Virus titre (plaque-forming units, PFU/mL), was determined by serially diluting the virus stock and counting the number of plaques in duplicate wells of MDCK cells containing between 10 and 100 plaques so that each plaque formed was due to one infective virus particle.

#### Separation of influenza virus from unbound gold AuNPs by filtration

WSN/H1N1 virus stock (10 µL) was mixed with 10 µL AuNPs and placed on a rocking platform for one hour at 37 °C. After this incubation, 980 µL PBS was added to the virus-AuNPs mixture. The mixture was then centrifuged on a 100 kDa cut off Nanosep filter for one minute at 10,000 g. Following resuspension of the residual ∼100 µL retentate in PBS, the cycle of washing by centrifugation and resuspension was repeated five times. The final retentate and the filtrates were then subjected to analysis by UV-Vis spectroscopy to quantify AuNPs and then by SDS-PAGE electrophoresis to identify proteins.

#### Pre-treatment of MDCK cells with peptides and functionalised AuNPs

The medium from confluent monolayers of MDCK cells was removed and 10 µL free peptide or AuNPs functionalised with different mole % of ligand, mixed with 440 µL of DMEM, was added and incubated at 37°C for one hour. The cell monolayers were then washed with different solutions, as indicated in figure legends and a viral plaque assay was performed to determine the antiviral activity of any cell-associated free peptide or AuNPs functionalised with different mole % of FluPep ligand.

#### Measurement of association of AuNPs with cells

MDCK cells were cultured in six-well plates as stated above for virus plaque assay. Cell monolayers were incubated for one hour at 37°C with AuNPs as described in the figure legends. The cell monolayers were washed twice with 3 mL PBS to remove any residual free AuNPs. Various washes and trypsinised cells, detailed in figure legends, were collected and their AuNP content quantified by measuring the absorption at 520 nm. In some experiments, the anti viral activity of the AuNPs that remained associated with the cells after the washes with PBS or 2 M NaCl was measured using the plaque assay.

#### Treatment of MDCK cells with glycosidases

Following fixation with 10% (v/v) neutral buffered formalin, cells were washed three times with PBS and then incubated with 2 mL of PBS containing 25 mM glycine pH 7.2 to block any remaining partially active fixative. The blocking medium was discarded after 15 min and cells were washed with three times with PBS. Cells were then incubated with the following enzymes:

i. To degrade heparan sulfate, 400 µL heparinase II and III (100 µL heparinase II and 200 µL heparinase III and 100 µL of 100 mM sodium acetate and 0.1 mM calcium acetate, pH 7.0 buffer; gifts from Dr. Edwin Yates, University of Liverpool) were added to cells;
ii. To degrade chondroitin sulfate (including dermatan sulfate): one mL of chondroitinase ABC (Sigma-Aldrich; 1.25 mU in PBS with x mg/mL bovine serum albumin) was added to the cells. These enzymes may contain traces of protease. Therefore, bovine serum albumin was added with these enzymes to ensures cell proteins were not degraded;
iii. To degrade sialic acid, neuraminidase 200 µL neuraminidase plus 800 µL 50 mM sodium acetate 4 mM CaCl_2_ 100 µg/mL BSA (pH 7.4) was added to cells.

In all cases, cells were incubated with the enzyme reaction mixtures overnight at 37°C.

#### SDS-PAGE

Samples (30 μL) for SDS-PAGE electrophoresis, were mixed with 10 μL 4xSDS-PAGE loading buffer [(50 % (v/v) glycerol, 10 % (w/v) SDS, 10 % (v/v) 2-mercaptoethanol in 0.3 M Tris-Cl (pH 6.8), coloured with bromophenol blue] and denatured by heating at 95°C for 10 min. Ten μL was then loaded onto a 12 % (w/v) SDS-polyacrylamide gel. Electrophoresis was done at 30 mA (per gel), 200 V for 50 min in running buffer [(50 mM Tris-Cl, 192 mM glycine and 0.1 % (w/v) SDS)].

After electrophoresis, the gels were incubated in fixative [(40 % (v/v) methanol, 10 % (v/v) acetic acid)] for 1 hour followed by two 5 min incubations in 10 % (v/v) methanol (5 min/each) and then washed in reverse osmosis water 3 times for 5 minutes. Gels were then either stained with Coomassie Blue (CBB R-250, ThermoFisher, UK) according to the manufacturer’s instruction or silver stained. Following incubation in 0.2 % (w/v) silver nitrate for 30 min, the gels were washed with water for 15 seconds and then dipped in freshly made developer solution [(2.5 % (w/v) Na_2_CO_3_, 0.03 % (v/v) formaldehyde)] until the colour of the solution turned brown. New developer was then used to further stain the gel until bands were stained to the required intensity. Stop solution (1 % v/v acetic acid) was added to stop the reaction. The gels were then washed with water six times for 5 minutes. Freshly made reducer (0.6 % w/v sodium thiosulphate, 0.3 % w/v potassium ferricyanide, 0.1 % w/v sodium carbonate) was used to remove excess silver and to clear the background. The gels were then quickly washed with a large volume of water. When required, gels were re-stained to increase the sensitivity of detection, starting by the addition of 0.2 % (v/v) silver nitrate for 30 min.

## Results and Discussion

### Binding of AuNP-FluPep ligand conjugates to virus

Previously, FluPep was suggested to exert its antiviral activity by binding to the influenza virus envelope, although these data were indirect, since they used an ELISA assay with wells coated with fluorescently labelled peptide by physisorption (Nicol et al., 2012) We took advantage of the exceptionally high extinction coefficient of AuNPs, which allows sensitive detection by UV-vis spectrophotometry (Haiss et al., 2007) to measure directly if FluPep bound to virus. This approach was made possible by the demonstration that FluPep functionalised AuNPs retain antiviral activity (Alghrair et al., 2019), and the substantial difference in size between the AuNP-FluPep ligand conjugate (diameter ∼8.8 nm) and the virus (diameter ∼100 nm). Virus and FluPep ligand conjugated AuNPs were incubated at 37°C for 30 minutes, as performed in the virus plaque assays (Alghrair et al., 2019). The FluPep-functionalised AuNPs were then separated from virus using a centrifugal filter with a cut-off of 100 kDa, which corresponds to around 10-12 nm. Measurement of the absorbance at 520 nm demonstrated that the gold AuNP-FluPep ligand conjugate was recovered in the filtrate and not in the retentate (Fig.1A). The presence of virus was determined by detection of viral proteins by SDS-PAGE. This demonstrated that the retentate, but not the filtrate, contained protein bands corresponding to viral proteins (Fig. 1B). The differences in the intensity of staining of the three major viral proteins, neuraminidase (NA), the viral proton channel (M2), and haemagglutinin (HA) likely arose from their relative numbers per virus particle and the known amino acid sequence-dependent difference in staining. Other weakly stained polypeptides may be due to cellular debris in the viral stock not being fully removed by centrifugation or to secreted cellular proteins. These data demonstrated that at the limit of detection of the assays, the FluPep ligand did not interact directly with the virus or did so very weakly, since virus and FluPep functionalised AuNPs were separated by the filtration step.

**Figure 1.**
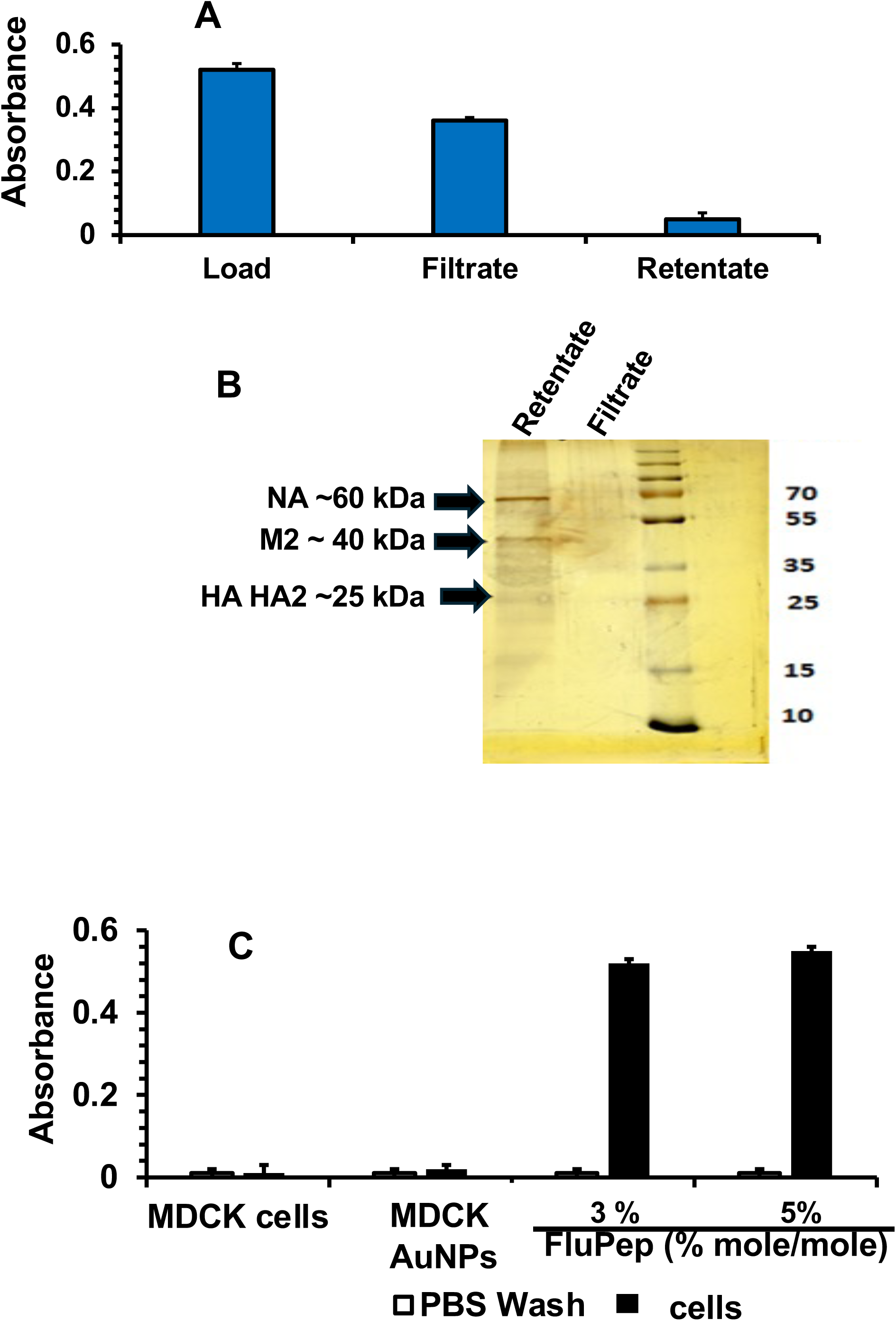
FluPep functionalised nanoparticles interact with MDCK cells but not with influenza virus. AuNPs functionalised with 5 % (mole/mole) FluPep were incubated with influenza virus for 1 h at 37 °C. Following dilution with PBS the mixture was separated by centrifugal filtration on a 100 kDa cut-off filter. (A) The absorbance at 450 nm of the retentate and the filtrate was measured to determine the presence of nanoparticles. Results are the mean ± SD of three experiments. (B) SDS-PAGE of the fractions from (A), the bands corresponding to major viral proteins are only present in the retentate and are indicated by arrows. (C) AuNPs and AUNPs functionalised with 5 % (mole/mole) FluPep were incubated with MDCK cells for 1 h at 37 °C. The DMEM containing unbound nanoparticles was removed and cells were then washed with 3 mL PBS and the absorbance in the PBS wash and the cells measured at 520 nm.

### Binding of AuNP-FluPep ligand conjugates to cells

To determine if the AuNP-FluPep ligand conjugate bound to MDCK cells, AuNP-FluPep were incubated at 37°C for 30 min with the conjugate in the same conditions as for the plaque assay (Alghrair et al., 2019). Following incubation with AuNPs, the cells were washed with PBS. The plasmon band maximum (520 nm) was used to determine the presence of cell-associated AuNPs as the samples had low background in this region. It should be noted that cells collected by scraping resulted in high scattering by cell clumps, so instead, cells were released by trypsin treatment. At 520 nm there was no detectable absorbance in the MDCK cells or in MDCK cells incubated with mix-matrix AuNPs. Thus, there was no detectable background signal from the cells and no detectable non-specific binding by the AuNPs passivated with the mix-matrix ligand shell (Fig. 1C), in agreement with previous work done on fibroblasts (Duchesne et al., 2012).

After incubation of MDCK cells with AuNPs functionalised with either 3% (mole/mole) or 5% (mole/mole) FluPep ligand, a substantial cell-associated absorbance at 520 nm, corresponding to the plasmon band of the AuNPs, was measured, which was resistant to washes with PBS (Fig. 1C). This was the ∼60% of the total AuNPs added to the cells. Thus, not only did the FluPep ligand conjugated AuNPs associate with the cells, but their cellular binding site(s) had a high capacity. Moreover, fixation of the cells with paraformaldehyde had little effect on the amount of FluPep ligand conjugated AuNPs bound to the cells (Fig. 2A).

**Figure 2.**
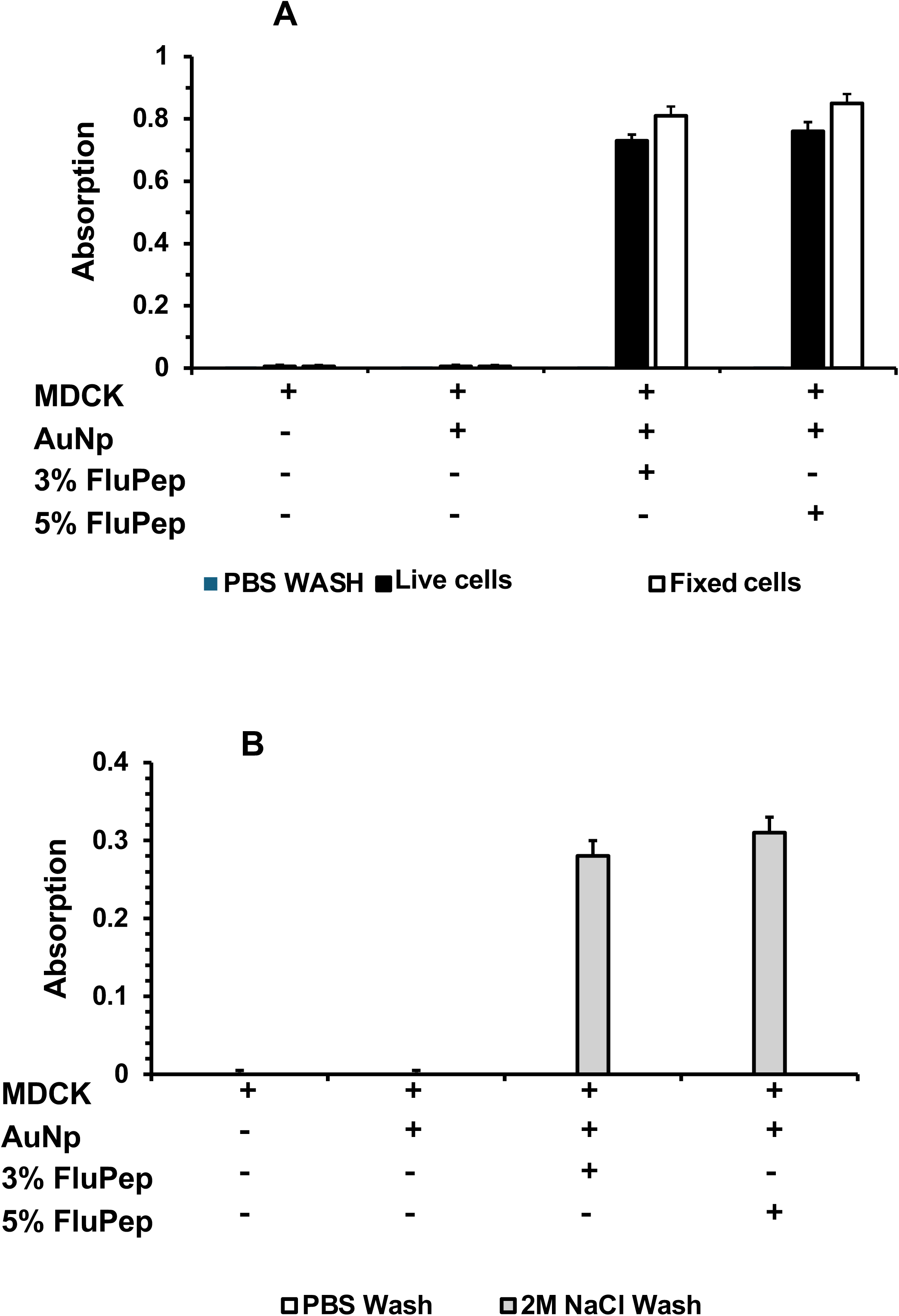
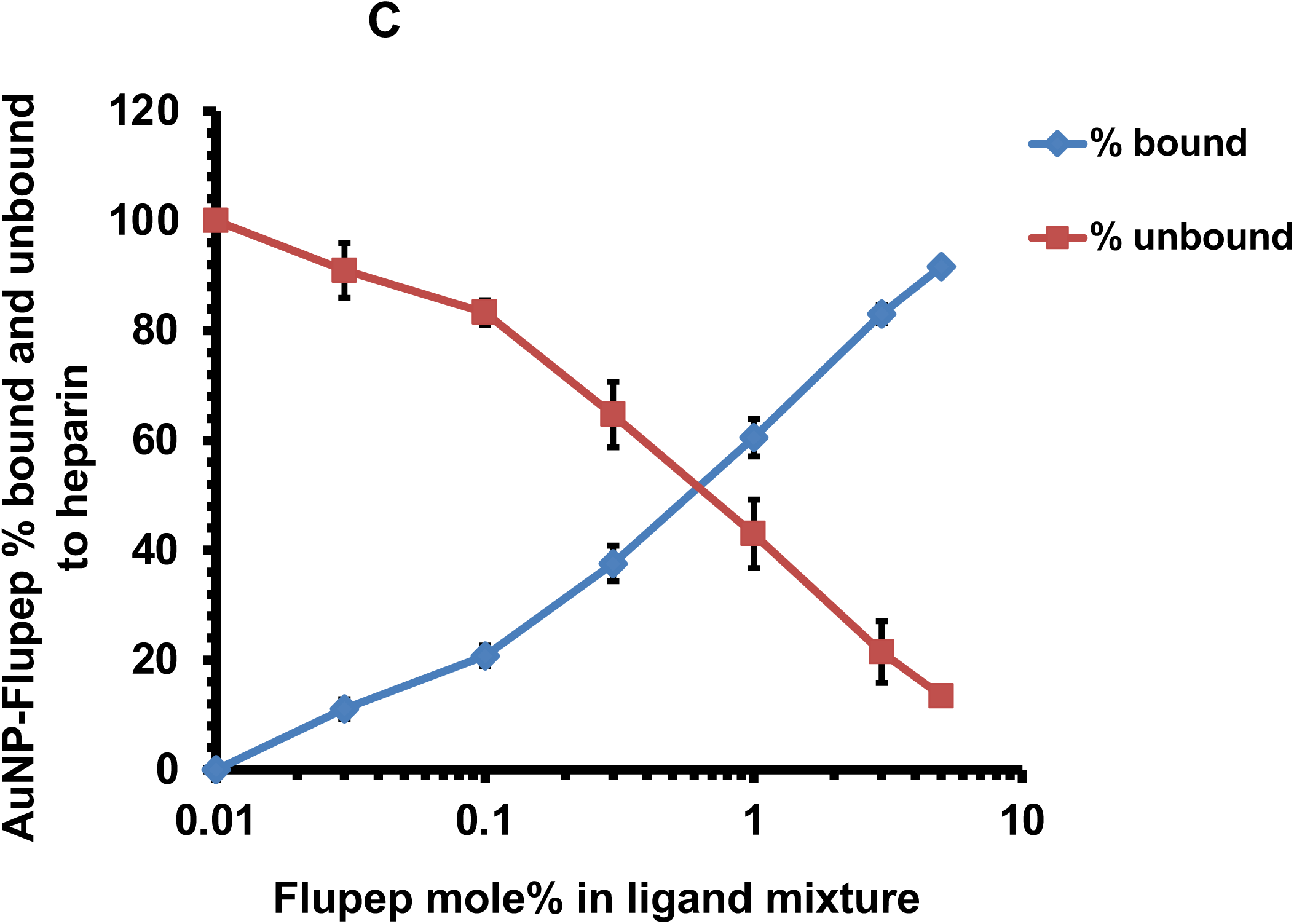
Interaction of FluPep functionalised AuNPs with MDCK cells and heparin. AuNPs functionalised with FluPep were incubated with live or fixed MDCK cells for 1 h at 37 °C in DMEM on a rocking platform. Then the DMEM with unbound AuNPs was removed and the cells were washed with 3 mL PBS and in some instances further washed with 3mL 2 M NaCl. The presence of AuNPs in the washes and in the cells after these were released by treatment with trypsin was determined by measuring the absorption at 2 nm. (A) Live and fixed MDCK cells incubated with AuNPs with just a mix-matrix (AuNPs) or functionalised with 3 % or 5 % (both mole/mole) FluPep ligand. B. Live MDCK cells incubated with AuNPs with just a mix-matrix (AuNPs) or functionalised with 3 % or 5 % (both mole/mole) FluPep ligand were sequentially washed with PBS and 2 M NaCl. Results are the mean ± SD (n=3). (C) AuNPs functionalised with different % (mole/mole) FluPep ligand were applied to a 100 µL heparin affinity column. The unbound fraction and following washing with PBS the fraction eluting with 2 M NaCl were collected and the AuNPs quantified by their absorption at 520 nm functionalised with different mole % FluPep ligand and washing the columns with PBS. The results are the mean±SD of three experiments.

A major class of high capacity binding sites on the cell surface and in the pericellular matrix are anionic polysaccharides, particularly the glycosaminoglycans. To determine whether the interaction of the AuNP-FluPep ligand conjugate with its cellular target(s) involved ionic bonding, cells were washed with PBS and then with 2M NaCl. The results showed that NaCl released over half of the cell bound AuNP-FluPep ligand conjugates (*c.f.* Figs 2A and 2B). The sulfated glycosaminoglycan HS represents a major class of strongly anionic molecules universally present on the surface of mammalian cells (Alotaibi et al., 2024). Heparin is a commonly used proxy for cell HS and thus, we tested whether the AuNP-FluPep ligand conjugated AuNPs would interact with a heparin affinity column. The results showed that AuNP-FluPep ligand conjugates, regardless of the mole/mole % functionalisation bound to a heparin affinity column. This binding was due to the FluPep ligand, since as the mole % of FluPep ligand increased, so did the percentage of gold AuNPs binding to the column (Fig. 2C).

### Binding of gold AuNP-FluPep ligand conjugates to fixed and enzyme-treated MDCK cells

The substantial binding of FluPep ligand-functionalised AuNPs to MDCK cells suggested that the peptide was interacting with abundant anionic components of the cell surface or pericellular matrix. The principal high-capacity anionic molecules in this environment are polysaccharides, particularly the glycosaminoglycans heparan sulfate (HS) and chondroitin sulfate (CS), together with sialic acid-containing glycans. Although negatively charged proteins are also present at the cell surface, they are generally of much lower abundance and charge density than these polysaccharides and are therefore less likely to account for the extensive binding observed. We, therefore, sought to determine which class of anionic glycans mediated the interaction of FluPep ligand-functionalised AuNPs with MDCK cells by selective enzymatic digestion.

To define better the anionic component(s) on the cells with which the gold AuNP-FluPep ligand conjugates interacted, and to understand whether this was simply an ion-exchange effect or due to selective affinity for particular species, cells were treated with enzymes that degrade three major classes of strongly anionic pericellular matrix molecules prior to measuring the binding of AuNPs. Since the binding of the AuNP-FluPep conjugates was similar in living and fixed cells (Fig. 2A), advantage was taken of the latter so as avoid the effect of the continuous biosynthesis of such molecules by cells (Duchesne et al., 2012, Sun et al., 2016).

Thus, a series of enzyme digestion experiments with neuraminidase (sialidase), heparinases and chondroitinase ABC were performed. Following a 24 h digestion of fixed cells, they were then incubated with AuNPs functionalised with 3% and 5% (both mole/mole) FluPep ligand for one hour at 37 ^°^C. In untreated cells, the absorption of the cell-associated AuNPs was between 0.47 and 0.52 (Fig. 3). In neuraminidase treated cells, the absorption was reduced by ∼70% to 0.13-0.14 (Fig. 3A). Similarly, treatment of the cells with heparinase reduced the amount of bound FluPep ligand functionalised gold AuNPs by ∼ 80%, whereas chondroitinase ABC digestion had no detectable effect (Fig. 3B). These data demonstrated that the FluPep ligand functionalised gold AuNPs bound to sialic acid-containing glycans and to heparan sulfate, the latter being consistent with the *in vitro* binding of the AuNPs to a heparin-affinity column (Fig. 2C). Interestingly, no chondroitinase ABC sensitive interaction was detected, indicating that the FluPep functionalised AuNPs had a degree of specificity for cellular anionic partners and consequently the interaction was not solely due to anion-exchange.

**Figure. 3.**
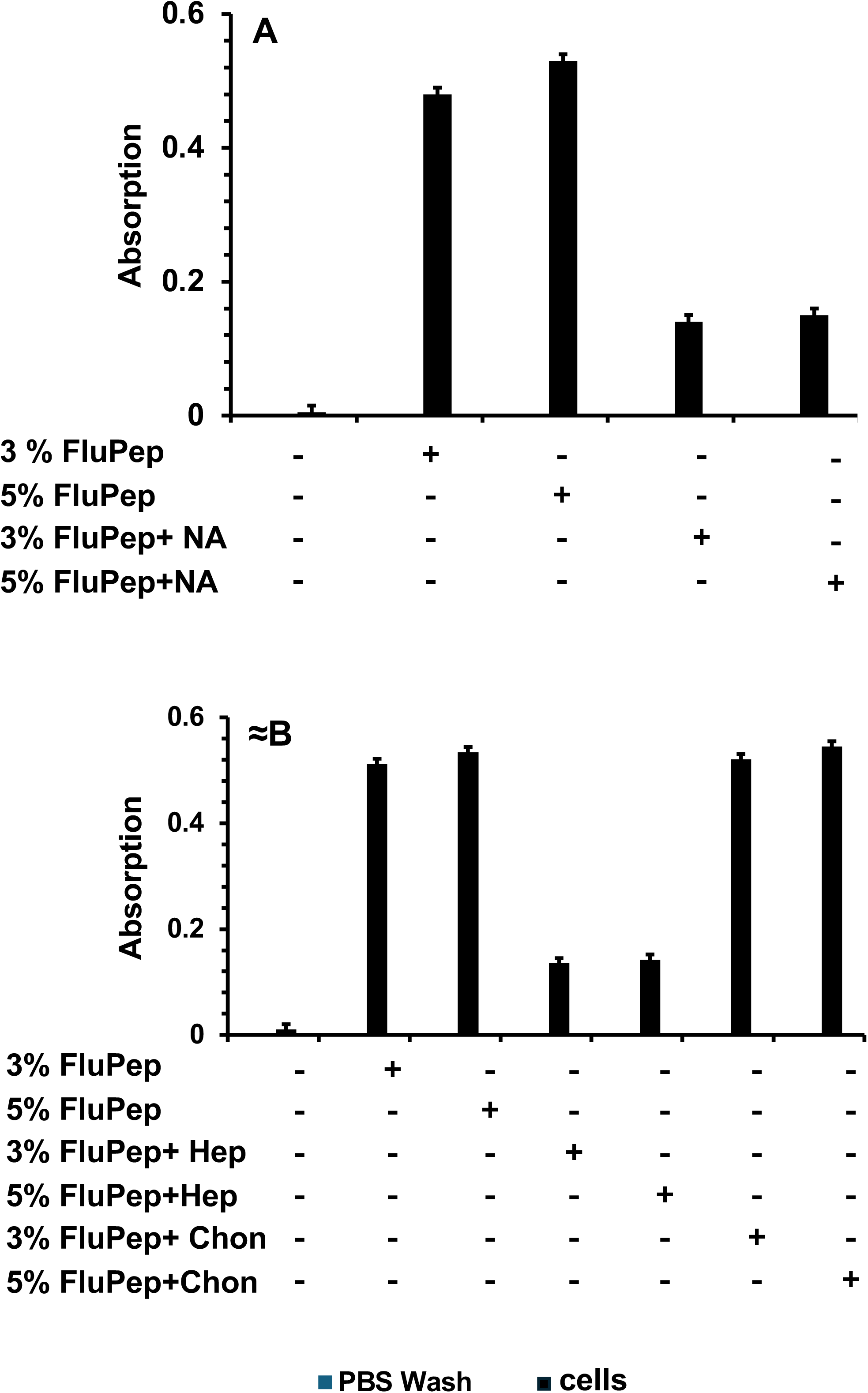
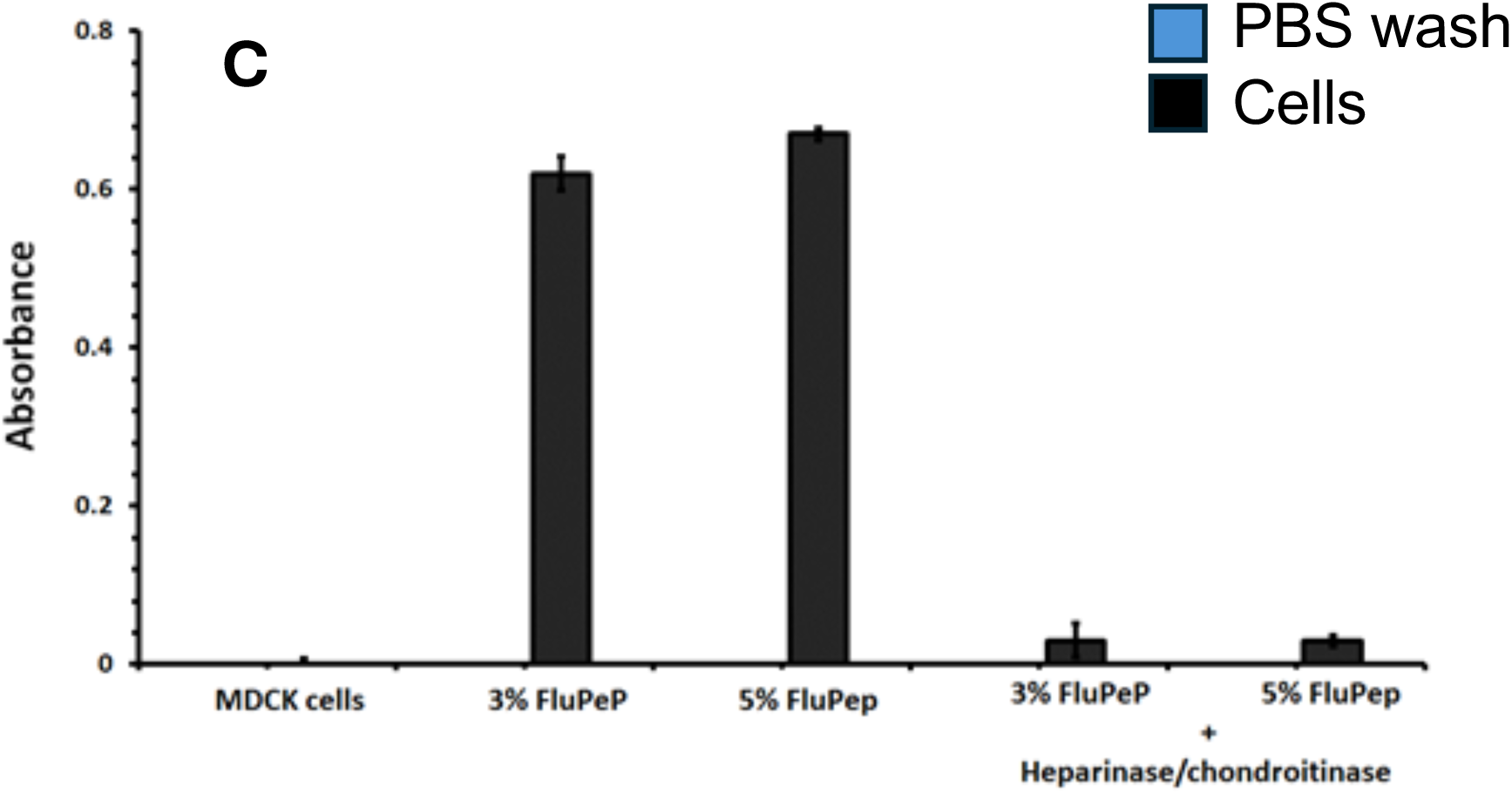
Sensitivity of binding of FluPep functionalised nanoparticles to degradation of polysaccharides on MDCK cells. Fixed MDCK cells were subjected to digestion with neuraminidase, heparinases or chondroitinase ABC overnight after which they were incubated with FluPep functionalised nanoparticles for 1 h on a rocking platform at 37 °C. After washing with PBS, cell-associated gold nanoparticles were measured by their absorption at 520 nm. For clarity the data have been divided into three panels, hence the controls are identical in each. (A) Neuraminidase (NA) treated MDCK cells, (B) heparinase (Hep) and chondroitinase ABC (Chon) treated cells, (C) Sequential treatment of cells with heparinases and chondroitinase (ABC). Results are the means±SD of three experiments.

Neuraminidase and heparinase treatment of fixed MDCK cells reduced binding of the FluPep ligand-functionalised gold AuNPs by well over half but did not abolish it. However, when cells were digested with both enzymes the amount of bound FluPep ligand functionalised gold AuNPs was reduced by ∼96% (Fig. 3C). These data indicated that sialic acid containing glycans and HS represent the major binding sites of FluPep ligand on MDCK cells. Digestion of one or the other of these polysaccharides is able to abolish most but not all of the binding of FluPep ligand functionalised gold AuNPs. Thus, it is likely that it is through these interactions that FluPep ligand exerts its antiviral effect, which would be by direct competition for these viral binding sites on the cell surface and in the pericellular matrix. This is supported by the observation that that exogenously added heparin was able to block the cellular infectivity of an influenza pseudovirus (Skidmore et al., 2015) and that the viral HA protein’s interactions with sialic acid is integral to viral infectivity (Chang et al., 2021)

### Anti-viral activity of cell bound gold AuNP-FluPep ligand conjugates

Over half of the AuNP-FluPep ligand conjugate added to cells were bound (Fig. 2). A similarly large fraction of HS-binding growth factors bind and are trapped in pericellular matrix, although they remain mobile within this and so able to activate their cellular receptors ((Alotaibi et al., 2024). Therefore, we tested the idea that AuNP-FluPep ligand conjugates bound to the pericellular matrix of cells in the absence of free AuNP-FluPep ligand conjugates in the bulk cell culture medium would protect these from viral infection. First, we determined whether washing the cells with 2 M NaCl affected their viability or the ability of influenza virus to subsequently infect them. Thus, MDCK cells were washed with 2M NaCl and then infected with the virus in the presence of increasing concentrations of free FluPep peptide. The results demonstrated the 2M NaCl wash had no effect on viral infectivity or the inhibitory activity of FluPep. (Fig. 4A), since the dose response to free FluPep peptide was identical whether the cells had been washed first with 2M NaCl or not (Fig. 4A). Cells were then incubated with AuNP-FluPep ligand conjugates functionalised with different % (mole/mole) FluPep ligand, and after binding they were washed with PBS to remove unbound AuNP-FluPep ligand conjugates. After this wash, the cells were subjected to the viral plaque assay. In this scenario the only AuNP-FluPep ligand conjugates present would be those associated with the anionic polysaccharides of the pericellular matrix. The AuNP-FluPep ligand conjugates that remained bound to cells after the PBS wash were capable of inhibiting the infectivity of influenza virus (Fig. 4B). In contrast, after binding AuNP-FluPep to MDCK cells and then washing with 2 M NaCl prior to the plaque assay there was negligible inhibition of viral infectivity (Fig. 4C). Thus, the fraction of AuNP-FluPep ligand conjugate that could be removed from the cells with 2 M NaCl and so which was interacting with anionic components of the cell surface appears responsible for much of the antiviral activity afforded by the AuNP-FluPep ligand conjugate.

**Figure 4.**
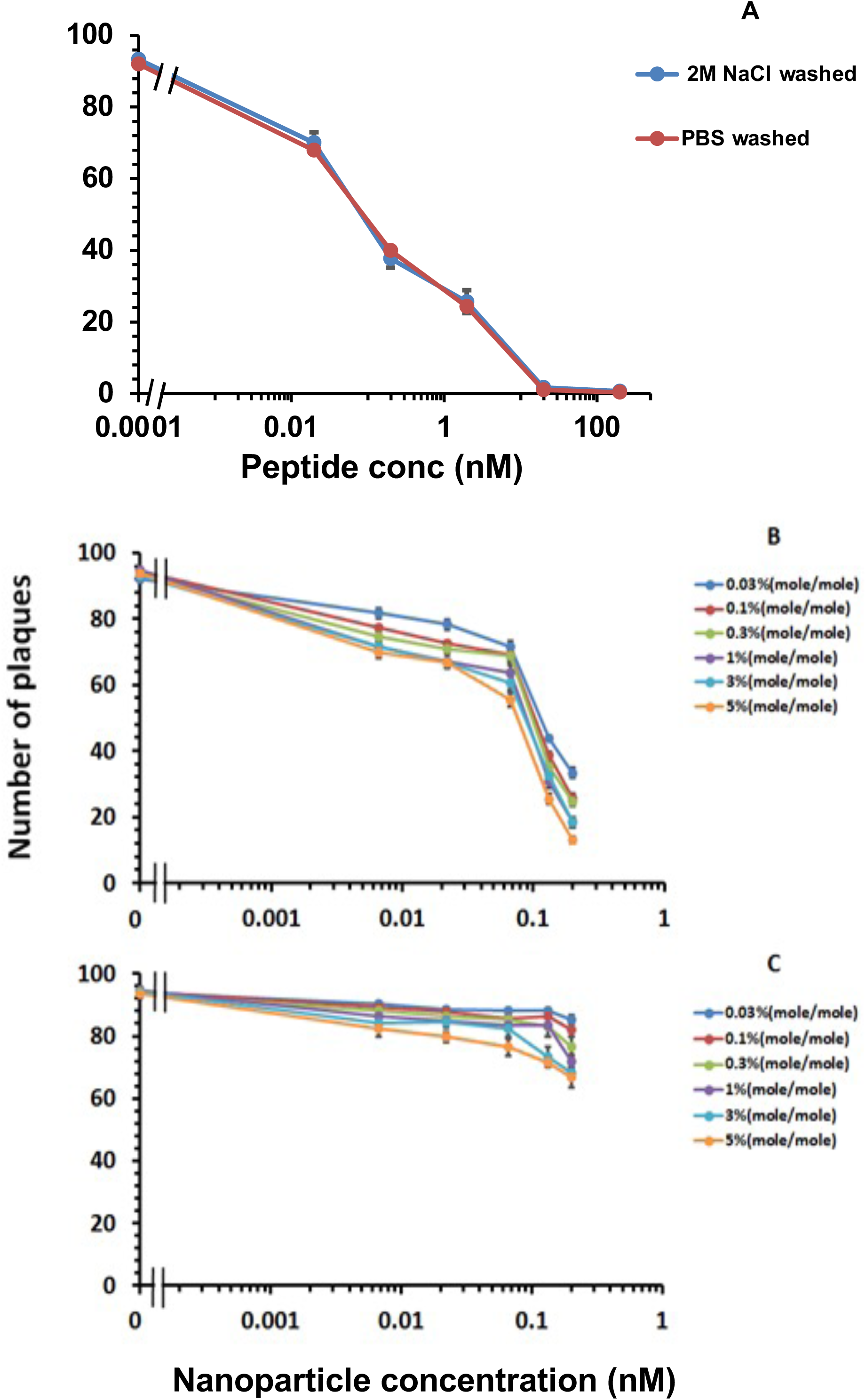
Anti-viral activity of cell bound gold AuNP-FluPep ligand conjugates. MDCK cells were subjected to different pre-trestments and then infected with influenza virus and a plaque assay performed. (A) A comparison of the anti-viral effect of FluPep ligand on cells washed with PBS or 2 M NaCl prior to the addition of FluPep and viral inoculum to determine if the short exposure to a hyper osmolar solution affected cell viability and so viral infectivity. (B, C) MDCK cells were incubated with AuNPs functionalised with different mole % FluPep and for 1 h at 37 °C on a rocking platform then washed with PBS (B) or 2 M NaCl (C), after which cells were infected with ‘flu virus and the number of plaques determined. Results are the mean ± SD (n=3).

### Contribution of the RRKK sequence in FluPep to its activity

In the original work, the sequence RRKK was added to either the N or C termini of the peptide derived from the sequence of cellular TKIP, in order to improve solubility in aqueous solution (Nicol et al., 2012). We reasoned that this basic tract of amino acids may be responsible, at least in part, for the interaction of FluPep ligand-conjugated AuNPs with anionic polysaccharides, which is an important component of FluPep’s antiviral activity (Figs 4B, C) and Table 2). Therefore, we next determined the contribution of the RRKK sequence to the antiviral activity of the peptides alone, of the peptide ligands (so incorporating the N-terminal sequence required for AuNP functionalisation) and of FluPep ligand-AuNP conjugates (Table 1). With the peptides alone, FluPep had the greatest antiviral activity and its IC_50_ was similar to that reported by Nicol et al (2012), while RRKKFluPep (so with the basic tract at the N-terminus, rather than the usual C-terminal position) possessed a comparatively lower antiviral activity and finally FluPepΔRRKK had the lowest antiviral activity of all (Fig. 5A Table 2). The antiviral activity of the corresponding peptide ligands was lower, in agreement with previous data that the N-terminal sequence CVVVTAA, required for conjugation to noble metal AuNPs, reduced antiviral activity of the free peptide (Alghrair et al., 2019). There was also a slight difference in the rank order of potency of these peptides: the position of the RRKK sequence (N- or C-terminal to the TKIP-derived sequence) had little effect, although the absence of this basic sequence again reduced the antiviral activity of the peptides (Fig. 5B, Table 2). The four basic residues on C-terminal to the matrix ligand (CVVVTAAARRKK) were without any antiviral effect (Fig. 5B).

**Figure 5.**
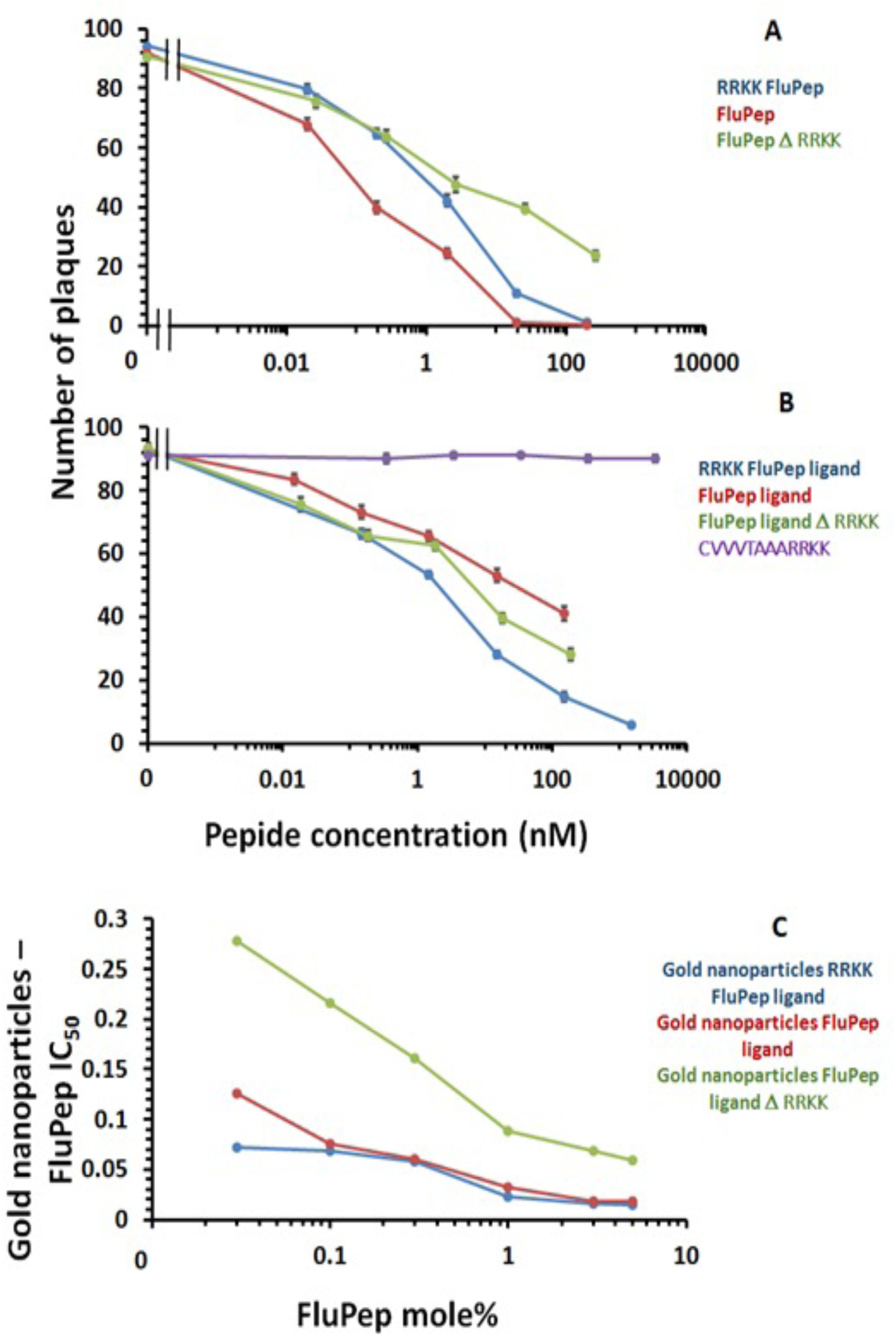
Influence of the RRKK sequence on the antiviral activity of FluPep. The anti viral activity of a range of variant FluPep peptides (Table 1) as well as the amino acid sequence comprising the comprising the peptidol, linker and the RRKK basic sequence (CVVVTAAARRKK) was tested in a plaque assay. (A) Antiviral activity of peptides alone, (B) antiviral activity of peptides in the form of ligands so with an N-terminal CVVVTAAA sequence, (C) antiviral activity of AuNPs conjugated to the peptides when at different mole %. Results are the mean ± SD (n=3).

**Table 2:**
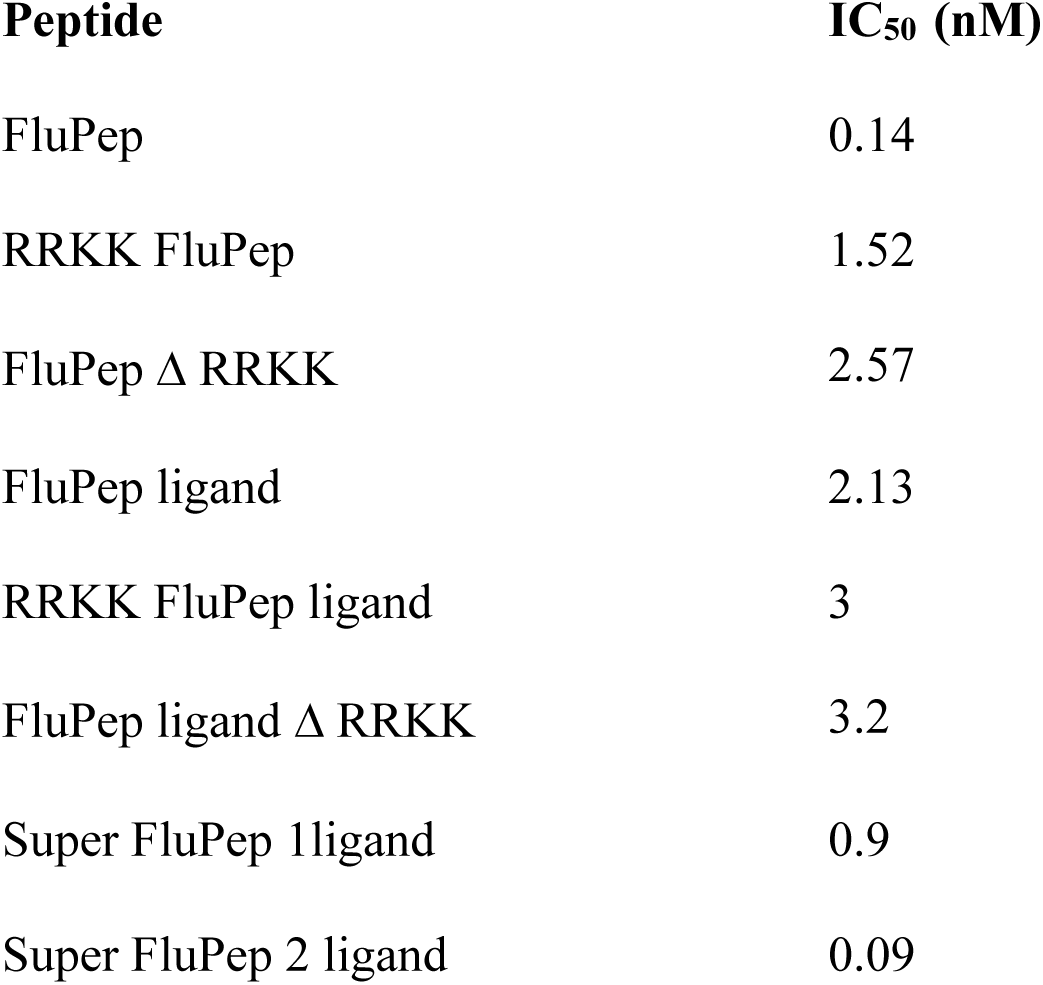
(IC_50_) of different free FluPep peptides in a plaque assay. The peptides are described in Table 1.

| Peptide | $IC_{50}$ (nM) |
| --- | --- |
| FluPep | 0.14 |
| RRKK FluPep | 1.52 |
| FluPep $\Delta$ RRKK | 2.57 |
| FluPep ligand | 2.13 |
| RRKK FluPep ligand | 3 |
| FluPep ligand $\Delta$ RRKK | 3.2 |
| Super FluPep 1ligand | 0.9 |
| Super FluPep 2 ligand | 0.09 |

When gold AuNPs were functionalised with these peptides, the position of the basic tetrapeptide, RRKK, made little difference to the antiviral activity of the AuNPs (Fig. 5C). However, as found for the free peptides, the absence of this basic sequence reduced the potency of the functionalised AuNPs considerably (Fig. 5C and Table 3). Thus, gold AuNPs functionalised with 3 % and 5% (both mole/mole) FluPepΔRRKK, possessed IC_50_ values of 278 pM and 59 pM AuNP-peptide conjugate, respectively (Fig. 5C), which are considerably higher than found for gold AuNPs functionalised with FluPep ligand.

### Design and activity of FluPep ligands with enhanced heparin;HS binding

FluPep ligand-functionalised gold AuNPs bound to heparin-affinity columns and to sialic acid and HS on cells (Figs 3A, B). Moreover, a variety of chemically modified heparin derivatives, some with very weak anticoagulant activities, have been shown to inhibit ‘flu virus H5N1 entry into cells using a pseudo virus assay (Skidmore et al., 2015). In the latter work, selectivity of the virus for particular sulfation patterns in heparin that would be more common in cellular HS was demonstrated by virtue of 6-*O* desulfated heparin possessing much less inhibitory activity than heparin itself, whereas 2-*O* desulfated heparin was considerably more active. There is a considerable amount of literature on protein structures and sequences that bind to heparin and HS (Ori et al., 2008, Ghezzi et al., 2017, reviewed Alotaibi et al., 2024). We reasoned that incorporating such sequences in place of the RRKK sequence might enhance the activity of the FluPep ligand further, by providing a higher affinity interaction with heparan sulfate. Indeed, such an approach has been used to increase the binding to HS and activity of growth factors (Martino et al., 2014). Consequently, two sequences were chosen. One was derived from placental growth factor 2 (PLGF2) and corresponds to a sequence used to enhance growth factor binding to HS (Martino et al., 2014). The second was derived from the canonical heparin binding site of fibroblast growth factor 10 (FGF10) (Li et al., 2016), but with the initial lysine residue of this sequence replaced by the corresponding arginine in FGF22 and the C-terminal T with the corresponding K from FGF7 (Xu et al., 2012) (Li et al., 2016) to increase affinity. These peptides are termed “super FluPep 1 ligand” and “super FluPep 2 ligand”, respectively (Table 1).

AuNPs functionalised with either of these peptides were as stable as those AuNPs functionalised with the original FluPep, in terms of their resistance to ligand exchange. The relative affinity for heparin of gold AuNPs functionalised with 0.1 % (mole/mole) FluPep ligand, Super FluPep 1 and 2 ligands was determined in terms of the concentration of NaCl required for elution from heparin affinity columns. Elution of FluPep ligand functionalised AuNPs was apparent at 0.6 M NaCl and 1 M NaCl (Fig. 6). In contrast, both super FluPep 1 ligand and super FluPep 2 ligand functionalised AuNP required 1.5 M to 2 M NaCl for elution, demonstrating that these two engineered constructs had a stronger electrostatic interaction with the polysaccharide (Fig. 6).

**Figure 6.**
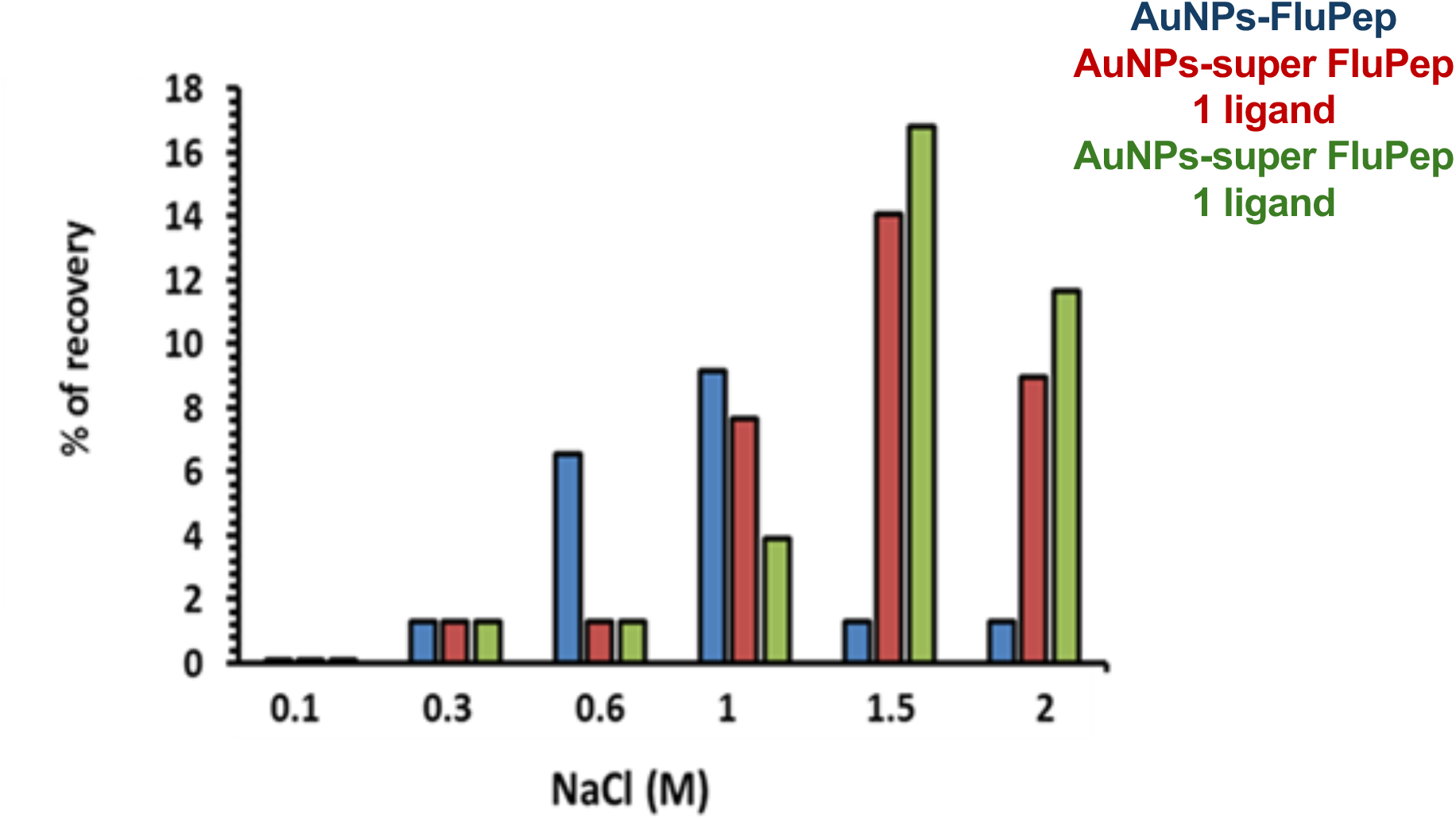
Comparison of concentration of NaCl required to elute gold AuNPs functionalised with FluPep, super Flupep1 and super Flupep2 from heparin affinity columns. The AuNPs functionalised with the respective FluPep ligand were applied to a heparin affinity column, which was then eluted with increasing concentrations of NaCl. AuNPs were quantified by their absorption at 450 nm. Results are the mean (n=2).

### Antiviral activity of super FluPep 1 and 2

The antiviral activity of the super FluPep 1 ligand and super FluPep 2 ligand (Table 1) alone and conjugated to AuNPs was measured. As noted previously (Alghrair et al., 2019) and observed in Figs 5A, B, the FluPep ligand which has the N-terminal CVVVTAA sequence necessary for incorporation into AuNP ligand shells is less potent than FluPep itself. Both super FluPep ligand peptides inhibited viral infectivity in a concentration-dependent manner and were more potent than FluPep ligand (Fig. 7). Thus, addition of the heparin binding sequences derived from PLGF and FGF10 rendered the respective FluPep ligand peptides as potent as the FluPep peptide and so more potent than FluPep ligand.

**Figure 7.**
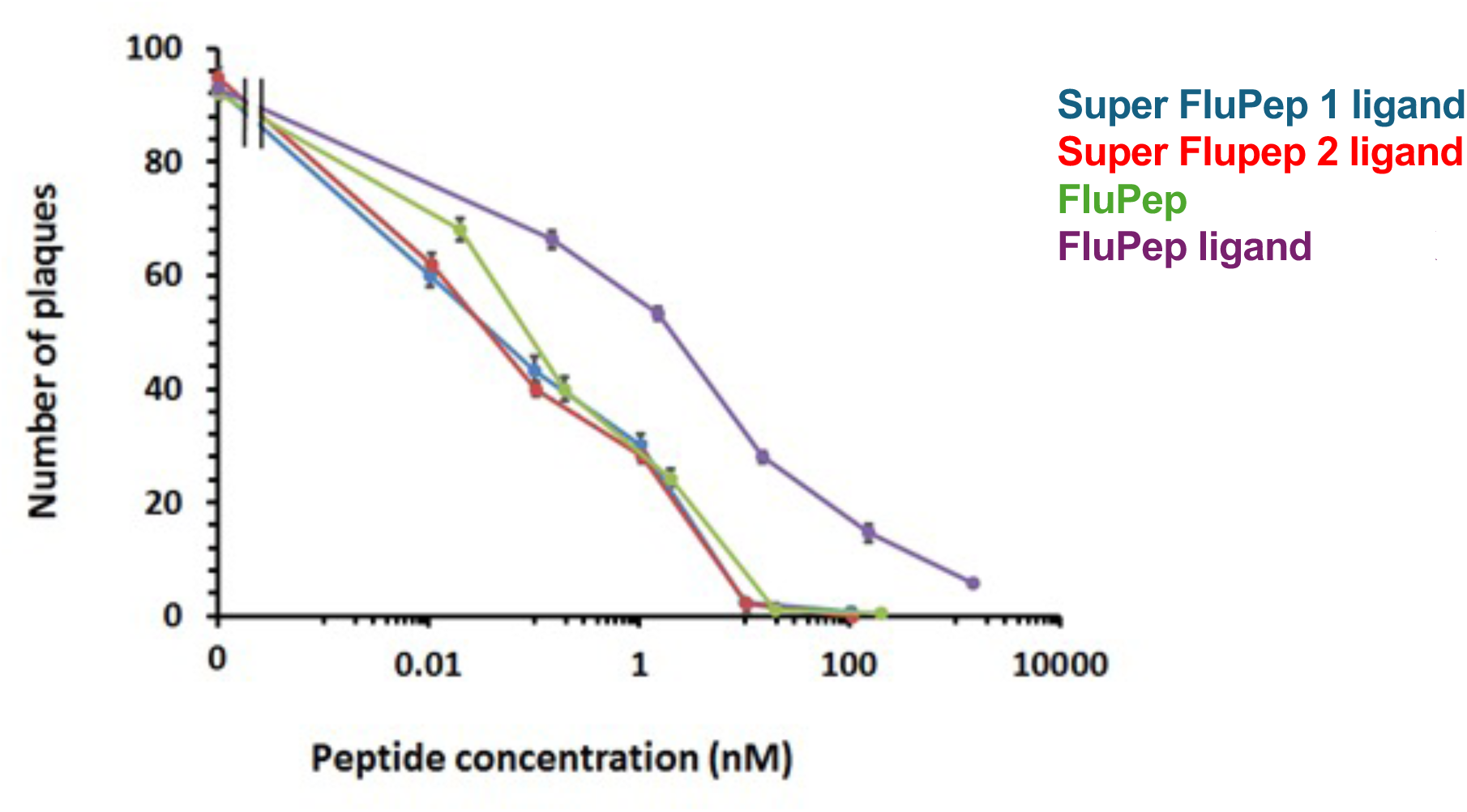
Anti-viral activity of TKIP peptides carrying various basic sequence extensions. Virus infection of a monolayer of MDCK cells was performed in the presence of FluPep, FluPep ligand, with the N-terminal extension necessary for conjugation to gold AuNPs, and two FluPep ligand peptides where the RRKK sequence of FluPep was replaced by a known heparin-binding sequence, Super FluPep 1 and Super FluPep2 (Table 1). Results are the mean ±SD (n=3).

The antiviral activity of AuNPs functionalised with different mole % of super FluPep 1 ligand and super FluPep 2 ligand was measured. The number of viral plaques was reduced as the concentration of gold AuNPs functionalised with either super FluPep 1 ligand or Super FluPep 2 ligand increased (Figs 8A,B). Thus, for gold AuNPs functionalised with 0.03% (mole/mole) Super FluPep 1 ligand and Super FluPep 2 ligand, the number of plaques started to decrease at 20 pM gold AuNPs and reached a minimum of around 13-17 plaques at 200 pM (Figs 8 A, B). As the grafting density of Super FluPep ligands was increased, so did the antiviral activity, to reach a maximum at 5 % (mole/mole) super FluPep1 and super FluPep2 ligands (Figs 8 A, B). It is noteworthy that the antiviral activity of Super FluPep1 ligand AuNPs and Super FluPep2 ligand AuNPs was greater than that of the free peptides consistent with the enhanced activity of FluPep conjugated to gold and silver AuNPs (Alghrair et al., 2019).

**Figure 8.**
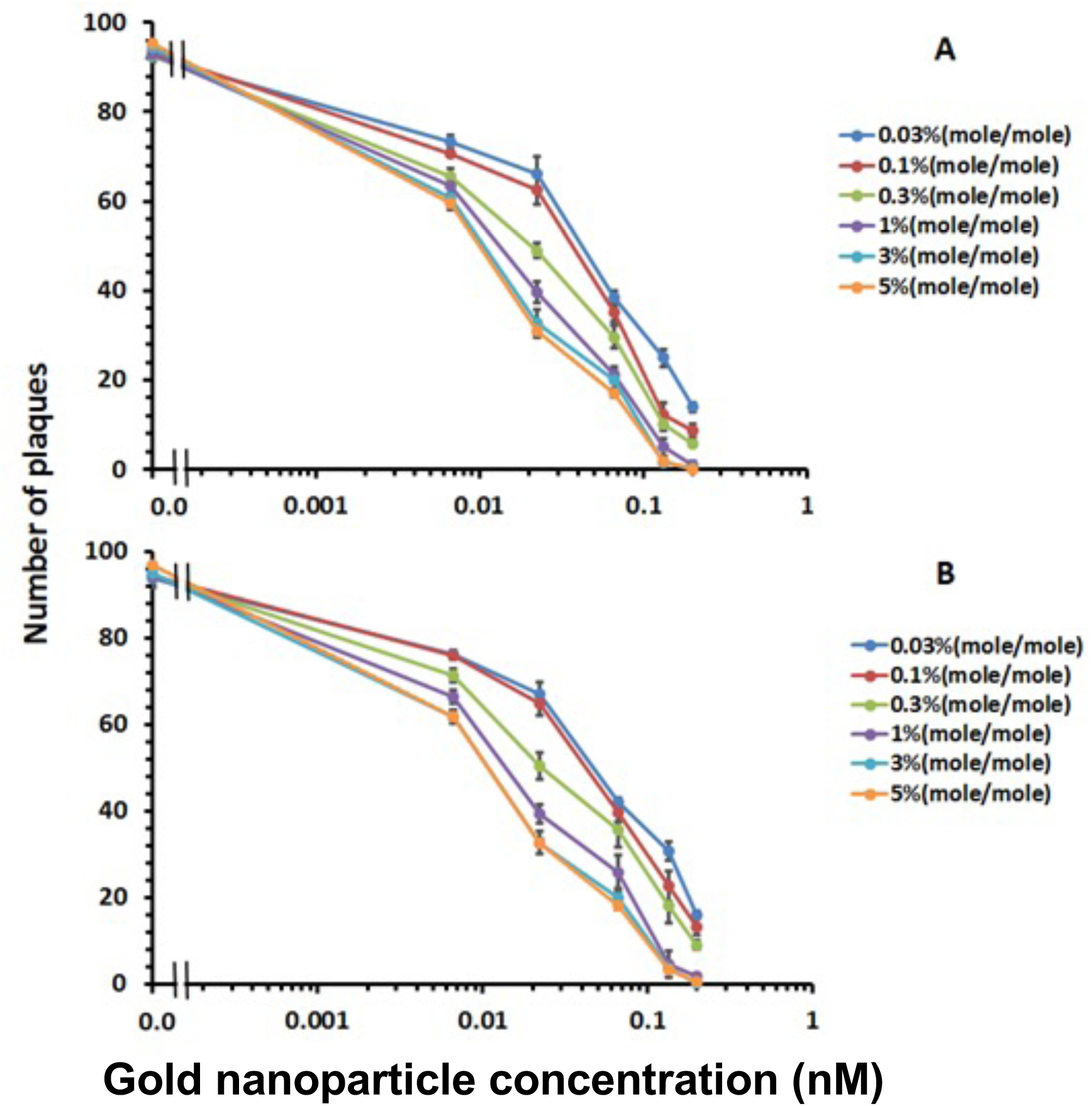
Anti-viral activity of gold AuNPs functionalised with super FluPep 1 and super FluPep 2. MDCK monolayers were incubated with influenza virus and AuNPs functionalised with different mole % of super FluPep 1 and super FluPep 2, following which the number of plaques was determined. Dependence of inhibition of plaque formation on the concentration of gold AuNPs and the % (mole/mole) of super FluPep 1 ligand (A) and super FluPep ligand 2 (B). Results are the mean ±SD (n=3).

## Conclusion

The FluPep peptide, when conjugated to noble metal AuNPs binds to anionic molecules in the cell pericellular matrix, rather than the influenza virus components. Interestingly, cell binding was somewhat selective, since FluPep ligand binding was sensitive to neuraminidase and heparinase, but not to chondroitinase ABC treatment, despite chondroitin sulfates possessing a high charge density. The RRKK basic tract (part of the FluPep amino-acid sequence) made an important contribution to the activity of FluPep ligand, whereas its replacement with amino acid sequences derived from the HS binding sites of proteins resulted in a substantially increased antiviral activity. Thus, cell binding is likely to be key to the mechanism of FluPep’s antiviral activity, since enhancing the FluPep peptide’s cell binding also enhanced its antiviral activity. The multivalent functionalisation of gold AuNPs also contributed to increased antiviral activity, as shown previously (Alghrair et al., 2019) and observed in Fig. 8. Therefore, these data demonstrate that engineering the sequence of FluPep with respect to its interactions with HS provides a means to a potent antiviral. As a number of other respiratory viruses, including RSV and SARS-CoV-2 also require interactions with cellular HS to infect host cells, AuNPs functionalised with heparan sulfate-binding amino acid sequences may be a route to new classes of anti respiratory viral drugs.

## Acknowledgements

Zaid K Alghrair was supported by a PhD studentship award from the Iraqi Ministry of Higher Education.

## Notes

### Competing Interest Statement

The authors have declared no competing interest.

